# LncRNA GUARDIN suppresses cellular senescence through a LRP130-PGC1α-FOXO4-p21-dependent signaling axis

**DOI:** 10.1101/695742

**Authors:** Xuedan Sun, Rick FrancisThorne, Xu Dong Zhang, Miao He, Shanshan Feng, Xiaoying Liu, Mian Wu

## Abstract

The long non-coding RNA GUARDIN functions to protect genome stability. Inhibiting GUARDIN expression can alter cell fate decisions towards senescence or apoptosis, but the underlying molecular signals are unknown. Here we show that GUARDIN is an essential component of a transcriptional repressor complex involving LRP130 and PGC1α which suppresses FOXO4 expression. GUARDIN acts as a scaffold to stabilize LRP130/PGC1α heterodimers and their occupancy at the FOXO4 promotor. Destabilizing this complex by silencing of GUARDIN, LRP130 or PGC1α leads to FOXO4-dependent upregulation of p21, thereby driving cells into senescence. We also found that GUARDIN expression was induced by rapamycin, a senolytic agent that suppresses cell senescence. FOS-Like Antigen 2 (FOSL2) acts as a transcriptional repressor of GUARDIN with increased levels in the presence of rapamycin resulting from downregulation of FOSL2. Together, these results demonstrate that GUARDIN inhibits p21-dependent senescence through a LRP130-PGC1α-FOXO4 signaling axis and moreover, GUARDIN contributes to the anti-senolytic activities of rapamycin.

## Introduction

Senescence is an irreversible state of cellular dormancy driven by various mechanisms such as replicative exhaustion resulting from telomere shortening or uncapping, oncogene activation, and genotoxic, nutrient and oxidative stress (1). These mechanisms commonly cumulate in the DNA damage response (DDR) leading to activation of the tumor suppressor p53 and the expression of cyclin-dependent kinase (CDK) inhibitors such as p21 and p16 that execute the senescent response. Although cellular senescence was initially regarded as a passive cell-autonomous anti-proliferation program echoing normal cellular aging (2, 3), it is increasingly appreciated that senescence plays an important role in many other physiological and pathological processes including tumor suppression, neurodegeneration and tissue remodeling (4, 5). Moreover, the senescence-associated secretory phenotype (SASP) resulting from changes in the secretome of senescent cells contributes to regulating inflammation, tissue microenvironment and age-related disorders (6–8).

As an important regulatory mechanism of cellular senescence that integrates a variety of extracellular and intracellular signals (1), the mechanistic target of rapamycin (mTOR; as known as mammalian target of rapamycin) plays an important role in regulating longevity (6). Inhibition of mTOR signaling by genetic or pharmacological approaches extends lifespan of various model organisms and mice with different genetic backgrounds (7–10). Indeed, treatment with the mTOR inhibitor rapamycin or its analogs (rapalogs) reduces cellular senescence (11–14). Nevertheless, the molecular mechanisms involved in rapamycin-mediated inhibition of senescence are not well understood.

An increasing number of noncoding RNAs have been found to be involved in regulation of cellular senescence (15). For example, the long noncoding RNA MIR31HG regulates the expression of p16 to modulate oncogene-induced senescence (16), whereas HOTAIR activates cellular senescence through p53-p21 signaling of the DDR (17). Moreover, the lncRNA OVAAL blocks cellular senescence through regulating the expression of the CDK inhibitor p27 (18). We have also found that the lncRNA GUARDIN plays an essential role in maintaining genomic stability (19). Silencing of GUARDIN was shown to induce cellular senescence although the mechanism responsible remains undefined.

In this report, we demonstrate that GUARDIN serves to facilitate assembly of the complex between LRP130/PGC1α that acts as repressor complex at the FOXO4 promoter. Silencing of GUARDIN disrupts this complex leading to FOXO4-dependent upregulation of p21, thereby driving cell entry into senescence. On the other hand, GUARDIN expression is upregulated by rapamycin treatment and this result from modulation of FOS-Like Antigen 2 (FOSL2) levels. FOSL2 normally transcriptionally represses GUARDIN but downregulation of FOSL2 by rapamycin releases this inhibition to promote GUARDIN expression. Thus, GUARDIN inhibits p21-dependent senescence induction through a signaling axis involving LRP130-PGC1α-FOXO4 moreover plays a functional role in the antagonistic actions of rapamycin on senescence.

## Results

### GUARDIN inhibits cellular senescence through suppressing transcriptional activation of p21

To validate the notion that GUARDIN protects cells from cellular senescence (19), we knocked down GUARDIN using shRNA in normal human adult foreskin fibroblasts (HAFF), non-small cell lung carcinoma cells (A549 and H1299), and hepatocellular carcinoma HepG2 cells, respectively (Figure EV 1A). We observed that shRNA silencing of GUARDIN induced senescence-associated-galactosidase (SA-gal) activity (Figure 1A), the formation of senescence associated heterochromatin foci (SAHF) as shown by increased levels of H3K9me3 (Figure 1B), and enhanced the senescence-associated secretory phenotype (SASP) as represented by elevated extracellular levels of IL-6 and IL-8 in A549 cells (Figure 1C) (20). Knockdown efficiencies varied from 50-90%, possibly due to the varied endogenous expression profiles of GUARDIN observed among the different cell lines (Figure EV 1A, B). Alternatively, overexpression of GUARDIN accelerated cell proliferation (Figure EV 1D, E) consistent with prior reports (19). Northern blotting analysis of GUARDIN in H1299 and A549 cells showed a single band of 985nt (Figure EV 1C), consistent with the three exon transcript reported previously (19). Thus, loss of GUARDIN induces senescence while its overexpression promotes proliferation in cell lines from different sources.

**Figure 1.**
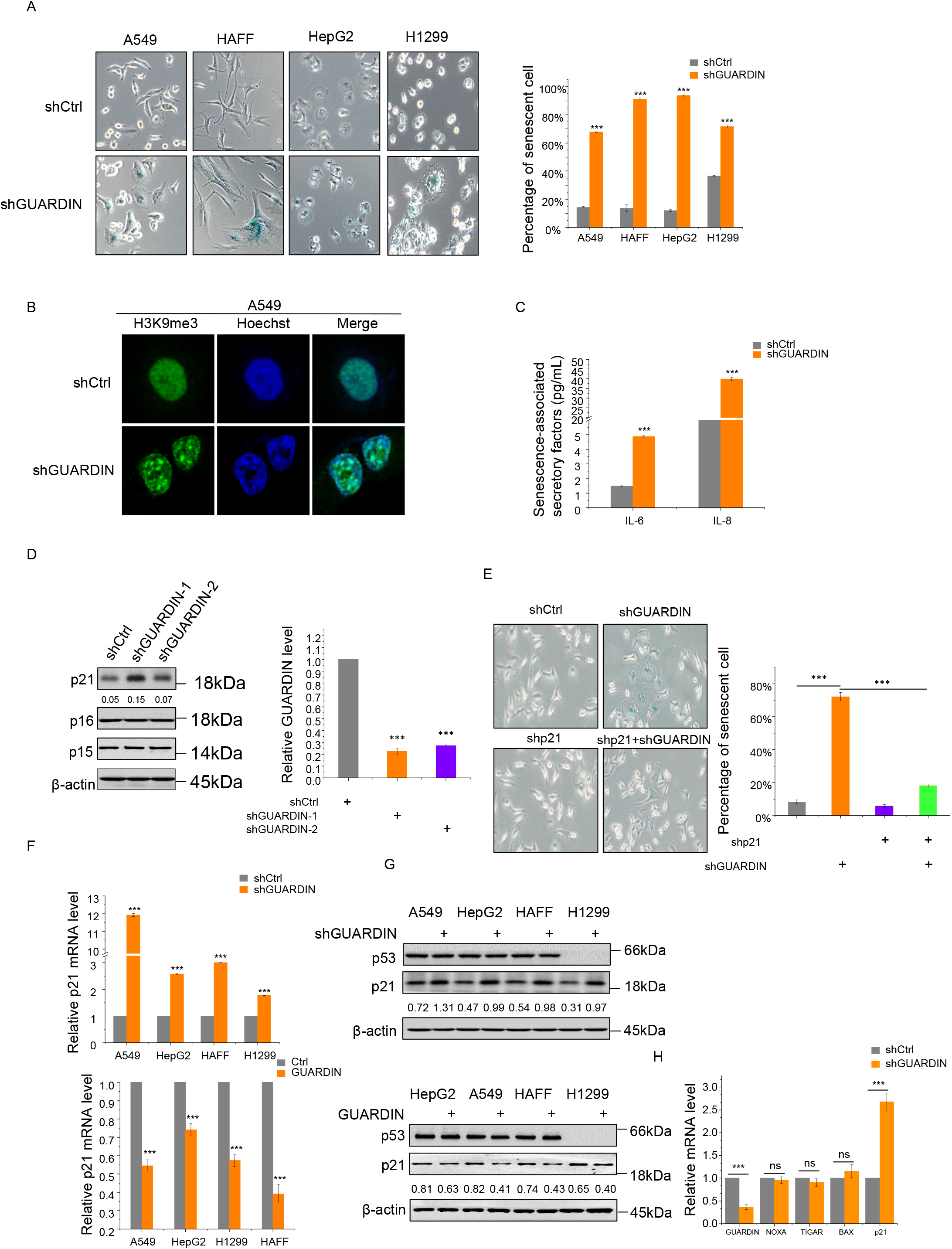
GUARDIN inhibits cellular senescence through suppression of p21. **A**. Senescence-associated β-galactosidase (SA-β-gal) staining in A549, HAFF, HepG2 and H1299 cells after 48h transduction with either negative control lentiviruses (shCtrl) or those targeting GUARDIN (shGUARDIN). Data shown as percentage of SA-β-gal-positive cells. **B**. Senescence-associated heterochromatin foci formation (SAHF) in A549 cells with either shCtrl or shGUARDIN. Representative IF staining with H3K9me3 antibodies (green) and nuclear Hoechst counterstaining (blue). **C.** Secretion of SASP factors IL-6 and IL-8 in A549 cells with either shCtrl or shGUARDIN. **D.** Western blotting for p21, p16 and p15 in A549 cells after 48h transduction with shCtrl or shGUARDIN-1, −2, respectively (left). Knockdown efficiencies analyzed by qPCR (right). **E.** SA-β-gal staining in A549 cells after 24h transduction with either shCtrl or shGUARDIN in combination with p21 shRNA (sh-p21). **F.** qPCR measurement of p21 mRNA in A549, HepG2, HAFF and H1299 cells with either GUARDIN shRNA knockdown (top) or GUARDIN overexpression (bottom). **G.** Western blotting for p53 and p21 in A549, HepG2, HAFF and H1299 cells with either GUARDIN shRNA knockdown (top) or GUARDIN overexpression (bottom). **H**. qPCR assays of NOXA, TIGAR, BAX and p21 mRNA levels in A549 cells with either shCtrl or shGUARDIN. (A, C, D-F, H) values are mean± s.e.m (n= 3). (A, C, F, H) two-tailed paired Student’s t test; (D) oneway ANOVA with Tukey’s multiple comparison post-test; (E) two-way ANOVA with Bonferroni’s multiple comparison post-test.

The induction of senescence following GUARDIN silencing was associated with upregulation of p21 protein, a factor which is well known to be involved in triggering senescence (21). In contrast, the expression of other senescence-inducing proteins, such as p15 and p16 (22), remained unaltered (Figure 1D). Indeed, p21 expression appeared critical for the induction of senescence caused by GUARDIN silencing, as co-silencing of p21 abolished SA-□-gal activity induced by GUARDIN knockdown (Figure 1E, Figure EV 1F). Together, these results suggest that GUARDIN silencing induces senescence through upregulation of p21. Indeed, we observed increased p21 mRNA levels in cells following knockdown of GUARDIN, and consistent with this notion, overexpression of GUARDIN caused a decrease in p21 transcription levels (Figure 1F). Of note, GUARDIN knockdown or overexpression did not affect the expression of p53 (Figure 1G), nor did it impact on p53 transcription activity, since knockdown of GUARDIN failed to influence the expression of other p53 transcription targets including Noxa, TIGAR and Bax (Figure 1H) (23). Moreover, GUARDIN knockdown also caused upregulation of p21 and activation of senescence in p53-null H1299 cells (Figure 1F, G). These data indicated that p53 is not involved in GUARDIN-mediated upregulation of p21.

To understand how silencing of GUARDIN could influence p21 expression, we first measured the effect of GUARDIN knockdown on the turnover of the p21 protein. Instructively there were no changes in the half-life time of p21 (Figure EV 1G) upon GUARDIN depletion, indicating that GUARDIN did not influence the post-translational stability of p21. Furthermore, treatment with the general transcription inhibitor actinomycin D (Act D) did not lead to noticeable changes in the turnover rate of p21 mRNA (Figure EV 1H) (24), suggesting that p21 upregulation was not a consequence of an increased p21 mRNA stability, rather, it occurred through upregulated transcription. Thus, GUARDIN appears to inhibit cellular senescence through repressing transcriptional activation of p21.

### GUARDIN facilitates LRP130/PGC1α interaction that mediates the transcriptional repression of p21

To investigate the mechanism responsible for GUARDIN-mediated suppression of p21, we interrogated the protein interactome of GUARDIN using RNA pulldown assays, and proteins that coprecipitated with biotin-labelled antisense probe against GUARDIN were identified by mass spectrometry (Figure 2A). Binding between GUARDIN and LRP130 was subsequently confirmed by Western blotting (Figure 2B). Consistently, the interaction of LRP130 and GUARDIN was readily detected using RNA immunoprecipitation (RIP) assay and RNA FISH (Figure 2C, D). Moreover, subcellular fractionation assays showed that GUARDIN and LRP130 were localized to both nuclear and the cytoplasmic compartments (Figure EV 2A, B), together indicating that LRP130 is a bona fide binding partner of GUARDIN.

**Figure 2.**
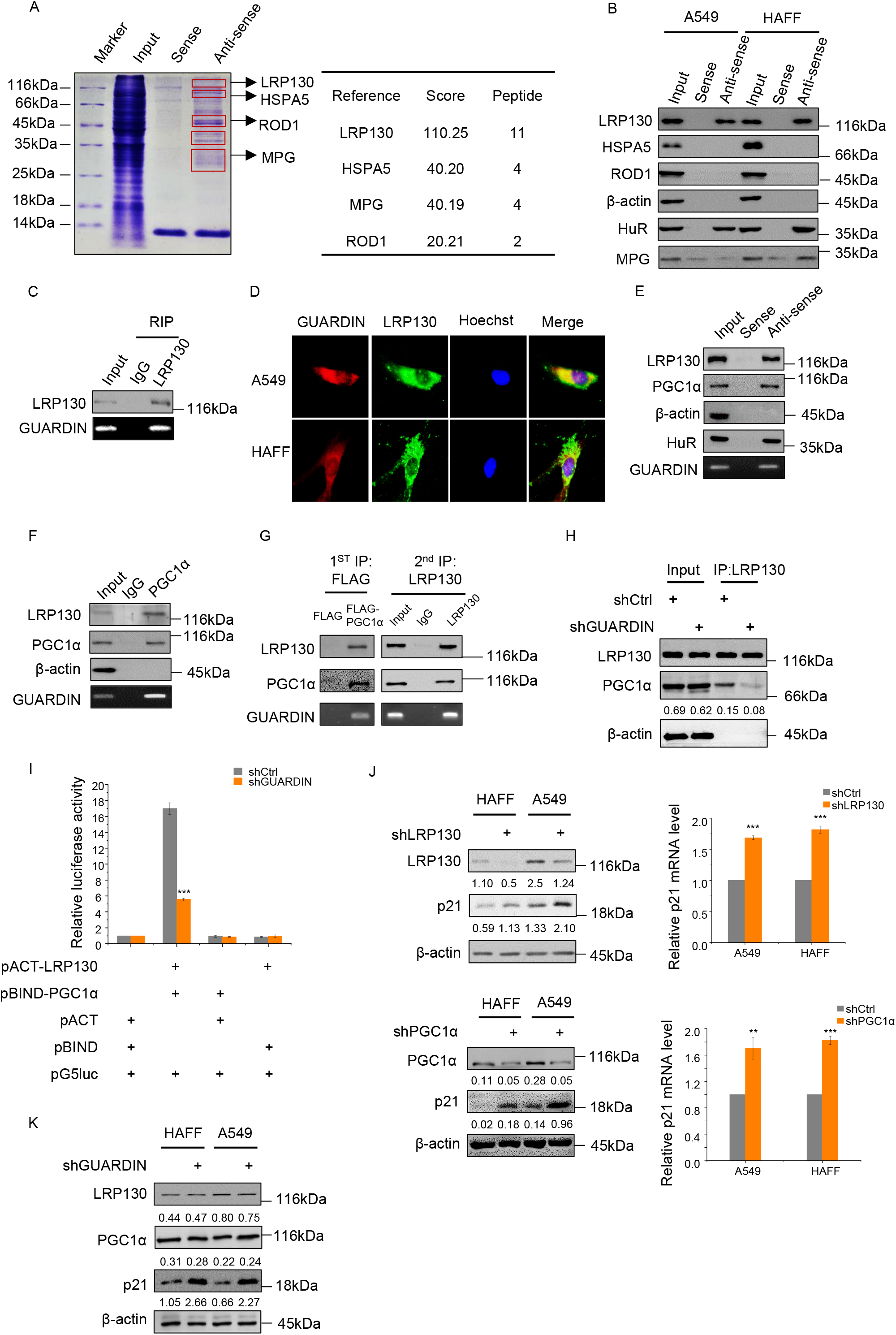
GUARDIN facilitates LRP130-PGC1α interaction which mediates transcriptional repression of p21. **A.** SDS-PAGE of RNA pulldown assays using biotin-labelled sense/antisense probes against GUARDIN from whole-cell lysates of A549 cells indicating putative GUARDIN-binding proteins (left); protein identities with high probabilities were determined by mass spectrometry (right). **B.** RNA pulldown assays interrogating putative GUARDIN-associated proteins identified in (A) from whole-cell lysates of A549 and HAFF cells. HuR and β-actin served as positive and negative controls, respectively. **C.** RNA immunoprecipitation (RIP) assays against IgG/LRP130 antibodies in whole-cell lysates of A549 cells. **D.** Subcellular localization of GUARDIN and its co-localization with LRP130. RNA FISH for GUARDIN (red) and IF for LRP130 (green) in A549 and HAFF cells. Nucleus were counterstained with Hoechst (blue). **E.** RNA pulldown assays using biotin-labelled sense/antisense probes against GUARDIN from wholecell lysates of A549 cells. GUARDIN levels were measured by RT-PCR and co-precipitated LRP130 and PGC1α detected by Western blotting. HuR and β-actin served as positive and negative controls, respectively. **F.** RIP assay using IgG/PGC1α antibodies from whole-cell lysates of A549 cells. GUARDIN, LRP130 and PGC1α levels were measured as per E. **G.** Two-step IP assays in whole-cell lysates of A549 cells transfected with FLAG-tagged PGC1α. First phase IPs were conducted with FLAG antibodies (left) and following elution with FLAG peptides, eluates were further subjected to second phase IPs with LRP130 antibodies (right). Samples were subjected to Western blotting and qPCR analysis for LRP130, PGC1α and GUARDIN, respectively. **H.** Co-immunoprecipitation (co-IP) between LRPl30 and PGC1α in A549 cells after 48h transduction with shCtrl or shGUARDIN. LRP130 was precipitated and samples subjected to Western blotting analysis for LRPl30, PGC1α and β-actin as loading control. **I.** Mammalian two-hybrid assays between pACT-LRP130 and pBIND-PGC1α in A549 cells after 48h transduction with shCtrl or shGUARDIN. Samples were subjected to the luciferase activity assays. **J.** LRPl30/PGC1α and p2l protein expression was measured by Western blot in A549 and HAFF cells after 48h transduction with shCtrl or shLRP130 (top left) or shPGC1α (bottom left) as indicated. qPCR assays for p21 mRNA levels were performed in parallel (right panels). **K.** Western blotting analysis of LRPl30, PGC1α and p2l protein levels in HAFF and A549 cells after 48h transduction with shCtrl or shGUARDIN. (I, J) values are mean± s.e.m (n= 3). (I, J) two-tailed paired Student’s t test.

LRP130 is a transcription factor shown previously to functionally complex with PGC1α to exert effects on gene expression (25). RNA pulldown assays performed with GUARDIN also showed the presence of PGC1α (Figure 2E) suggesting that GUARDIN may associate with the LRP130-PGC1α complex. Consistently, RNA immunoprecipitation (RIP) assay showed association between GUARDIN and PGC1α (Figure 2F). To verify that GUARDIN associates with the LRP130-PGC1α complex, we introduced FLAG-tagged LRP130 into A549 cells and conducted two-step IP assays from total protein extracts. In the first phase IP, FLAG antibodies precipitated LRP130 along with PGC1α and GUARDIN while in the second phase IP, PGC1α □ antibodies co-precipitated LRP130 and GUARDIN (Figure 2G), indicating GUARDIN, LRP130 and PGC1α exist as a ternary complex. Instructively, GUARDIN knockdown diminished the relative amount of LRP130 associated with PGC1α as shown in co-immunoprecipitation (Co-IP) and mammalian two-hybrid assays (Figure 2H, I), indicating that GUARDIN functions to facilitate interactions between LRP130 and PGC1α.

We also investigated the structural basis of the interaction of GUARDIN with LRP130 and PGC1α by using deletion-mapping experiments. Binding assays with in vitro-transcribed GUARDIN along with truncation mutants demonstrated that deletion of exon 3 (E3; −288/-985) diminished the binding of GUARDIN to LRP130 and PGC1α (Figure EV 2C). These results support the notion that E3 region of GUARDIN is the binding unit responsible for the association with LRP130 and PGC1α. Deletion mapping experiments with truncated mutants of LRP130 and PGC1α revealed that the N-terminal and C-terminal regions of LRP130 are required for its association with GUARDIN, whereas the C-terminus and central region of PGC1α are responsible for binding to GUARDIN (Figure EV 2D).

We next examined whether LRP130 and PGC1α play roles in GUARDIN-mediated repression of p21. SliRNA knockdown of either LRP130 or PGCla□resulted in upregulation of p21 at both mRNA and protein levels (Figure 2J), recapitulating the effects of GUARDIN depletion on p21 (Figure 2K). Thus, the association of LRP130 and PGC1α facilitated by GUARDIN appears necessary for GUARDIN-mediated repression of p21 expression.

### GUARDIN/LRP130/PGC1α represses transcription of FOXO4 gene

To investigate whether LRP130/PGC1α was involved directly in transcriptional regulation of p21, we interrogated the p21 promoter region for the presence of LRP13/PGC1α□binding sites. Indeed, using the UCSC browser predicted LRP130/PGC1α binding sites were found embedded within DNase I hypersensitive regions (DHSs) within the p21 gene promoter (−1870/−1701, −1430/-1221, −395/-1 upstream of the transcription start site) (Figure 3A, lower part). DHSs mark diverse classes of cis-regulatory regions, such as promoters and enhancers and their identification represents a powerful method of identifying the location of gene regulatory elements, including promoters (26–28). Nevertheless, neither LRP130 nor PGC1α were found to directly bind to p21 promoter using ChIP assays whereas TP53, a bona fide p21 transcriptional driver, was shown to bind to the p1(−1870/−1701) and p4 (−395/−200) binding sites (Figure 3A, upper part). These results implied that the upregulation of p21 by LRP13/PGC1α was unlikely to be mediated through direct transcription.

**Figure 3.**
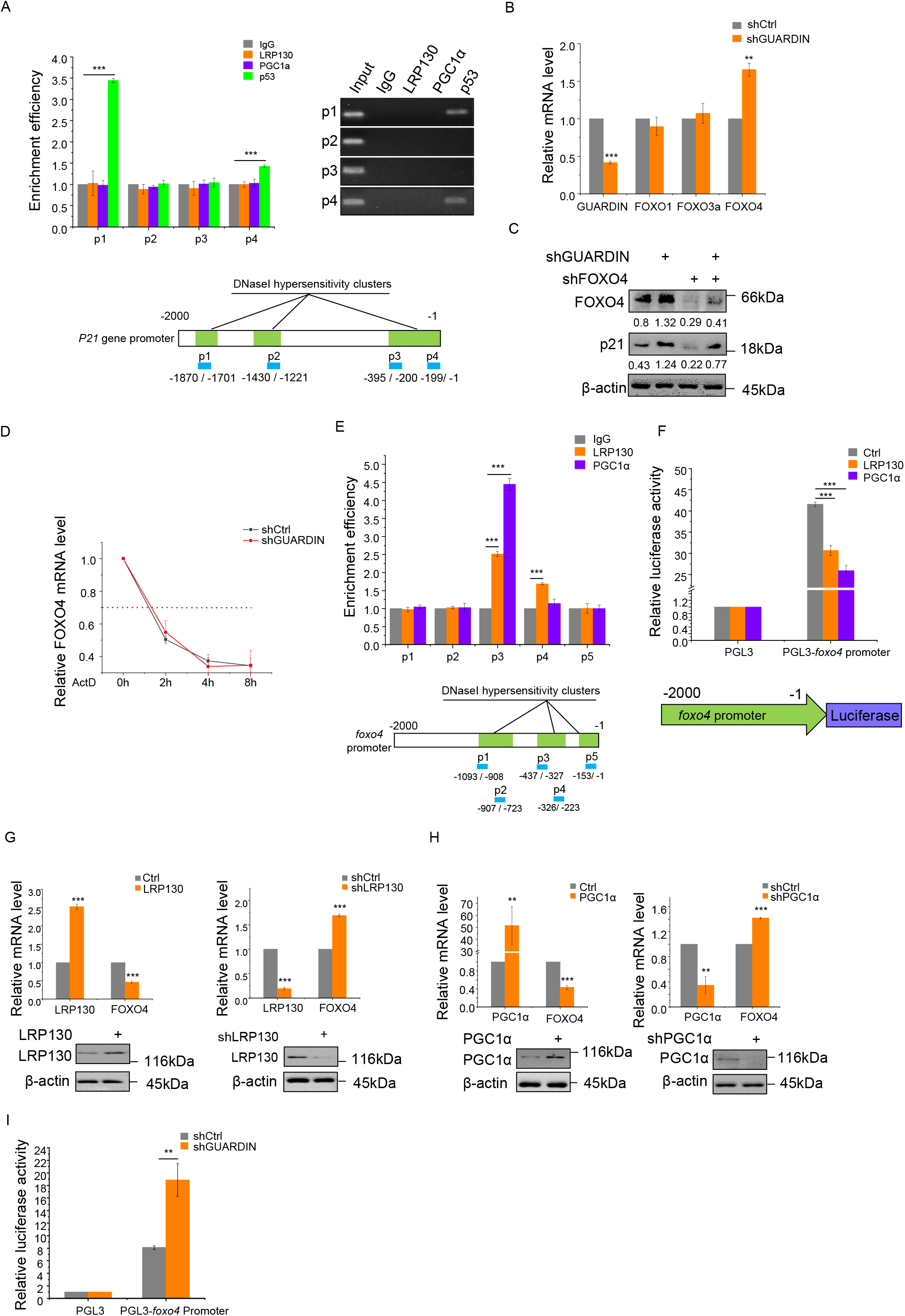
LRP130/PGC1α negatively regulates FOXO4 transcription. **A.** ChIP assays detecting binding of LRPl30/PGC1α to the p2l promoter using qPCR and RT-PCR (top left and right, respectively). IgG and p53 served as a negative and positive controls, respectively. Schematic illustrations of the putative LRPl30/PGC1α binding sites within DNase I hypersensitive regions of the p21 promoter (bottom). **B.** qPCR assays for GUARDIN, FOXO1, FOXO3a and FOXO4 in A549 cells after 24h transduction with shCtrl or shGUARDIN. **C.** Western blotting assays for FOXO4 and p21 protein expression in A549 cells after 48h transduction with shCtrl or shGUARDIN alone or in combination with shFOXO4. β-actin served as loading control. **D.** Half-life times of FOXO4 mRNA in A549 cells with shCtrl or shGUARDIN measured by qPCR after treating cells with 1 mg/ml of actinomycin (ActD) for the indicated times. **E.** ChIP assays detecting binding of LRPl30/PGC1α to putative binding sites in the FOXO4 promoter using qPCR (top). Data were normalized to the IgG negative control. Schematic illustration of the LRPl30/PGC1α binding sites within DNase I hypersensitive regions of the FOXO4 promoter (bottom). **F.** Luciferase assays conducted in A549 cells transfected with LRPl30 or PGC1α using the pGL3 (negative control) or pGL3-FOXO4 promoter reporter plasmids. **G.** qPCR assays for LRP130 and FOXO4 mRNA levels (upper) and Western blotting analysis for LRP130 protein level (lower) in A549 cells after 48h transduction with LRP130 overexpression or knockdown. **H.** qPCR assays for PGC1α and FOXO4 mRNA level (upper) and Western blotting analysis for PGC1α protein level (lower) in A549 cells after 48h transduction with PGC1α overexpression or knockdown. **I.** Luciferase activity assays in A549 cells transduced with shCtrl or shGUARDIN using the pGL3 (negative control) or pGL3-FOXO4 promoter reporter plasmids. (A, B, D-I) values are mean± s.e.m (n= 3). (B, G-I) two-tailed paired Student’s t test; (A, E, F) oneway ANOVA with Tukey’s multiple comparison post-test.

To clarify the mechanism by which GUARDIN suppresses p21 expression and cellular senescence, we carried out comparative RNA sequencing (RNA-seq) analysis for differe-ntially expressed transcripts in A549 cells with and without GUARDIN shRNA knockdown (Figure EV 3A). Notably the p21 promoter region is known to contain consensus forkhead binding elements (29) with the RNAseq analysis revealing significant association with FoxO signaling through enrichment of the forkhead box proteins, FOXO1, FOXO3a and FOXO4 in GUARDIN knockdown cells (Figure EV 3B, Dataset EV2). Verification using qPCR showed that FOXO4, but not FOXO1 and FOXO3a mRNA levels were increased upon GUARDIN knockdown (Figure 3B). Importantly, knockdown of FOXO4 not only reduced p21 expression but also diminished the increase in p21 caused by GUARDIN knockdown (Figure 3C). Collectively, these results suggest that GUARDIN-mediated repression of p21 may be due to suppression of FOXO4. Silencing of GUARDIN had no effect on the turnover rate of FOXO4 mRNA when treated with ActD (Figure 3D), suggesting that regulation of FOXO4 by GUARDIN was unlikely to affect FOXO4 mRNA stability. In support of this notion, we identified multiple consensus LRP130 and PGC1α□binding sites in the DNase I hypersensitive region (−1093/−723, −437/−223, −153/−1) of the FOXO4 gene promoter (Figure 3E, bottom). ChIP assays showed that LRP130 and PGC1α□bound to the P3 and to a lesser extent the P4 regions (−437/−327 and −326/−233, respectively; Figure 3E, top).

We then tested the functional capacity of LRP130 and PGC1α□to interact with the F0X04 promoter using luciferase reporter constructs containing the FOXO4 promoter with or without intact LRP130/PGC1α binding sites (Figure 3F, bottom). Reporter assays conducted in 293T cells showed that the transcriptional activity of the FOXO4 reporter construct with intact LRP130/PGC1α binding sites was lower than that of the control (Figure 3F). Consistently, shRNA knockdown of LRP130 or PGC1α resulted in upregulation of FOXO4 mRNA expression, while the overexpression of LRP130 or PGC1α downregulated FOXO4 mRNA levels (Figure 3G, H). Silencing of GUARDIN increased the transactivation of FOXO4 promoter as shown by luciferase assay (Figure 3I). Taken together, these findings established that GUARDIN suppresses p21 expression through facilitating LRP130/PGC1α-mediated transcriptional repression of FOXO4.

### GUARDIN is transcribed by FOSL2 that is responsive to rapamycin

The senolytic agent rapamycin inhibits senescence through a number of different mechanisms including but not only, inhibition of mTOR (11, 12). We investigated whether GUARDIN contributes to or even cooperates with rapamycin to inhibit senescence. Indeed, rapamycin treatment resulted in upregulation of GUARDIN, both in a time- and dose-dependent manner (Figure 4A, B). Intriguingly, the downregulation of p21 by rapamycin treatment was mediated through GUARDIN, since the p21-dependent cellular senescence caused by GUARDIN depletion was reversed by rapamycin treatment (Figure 4C, D). Rapamycin treatment increased the relative amount of LRP130 associated with PGC1α as shown in mammalian two-hybrid assays (Figure 4E), recapitulating the effects of GUARDIN. Therefore, GUARDIN is not only a rapamycin-responsive lncRNA, but also it contributes to the rapamycin-mediated inhibition of senescence. Moreover, the levels of p53 did not change upon rapamycin treatment (Figure 4C), implying that rapamycin does not regulate GUARDIN through p53.

**Figure 4.**
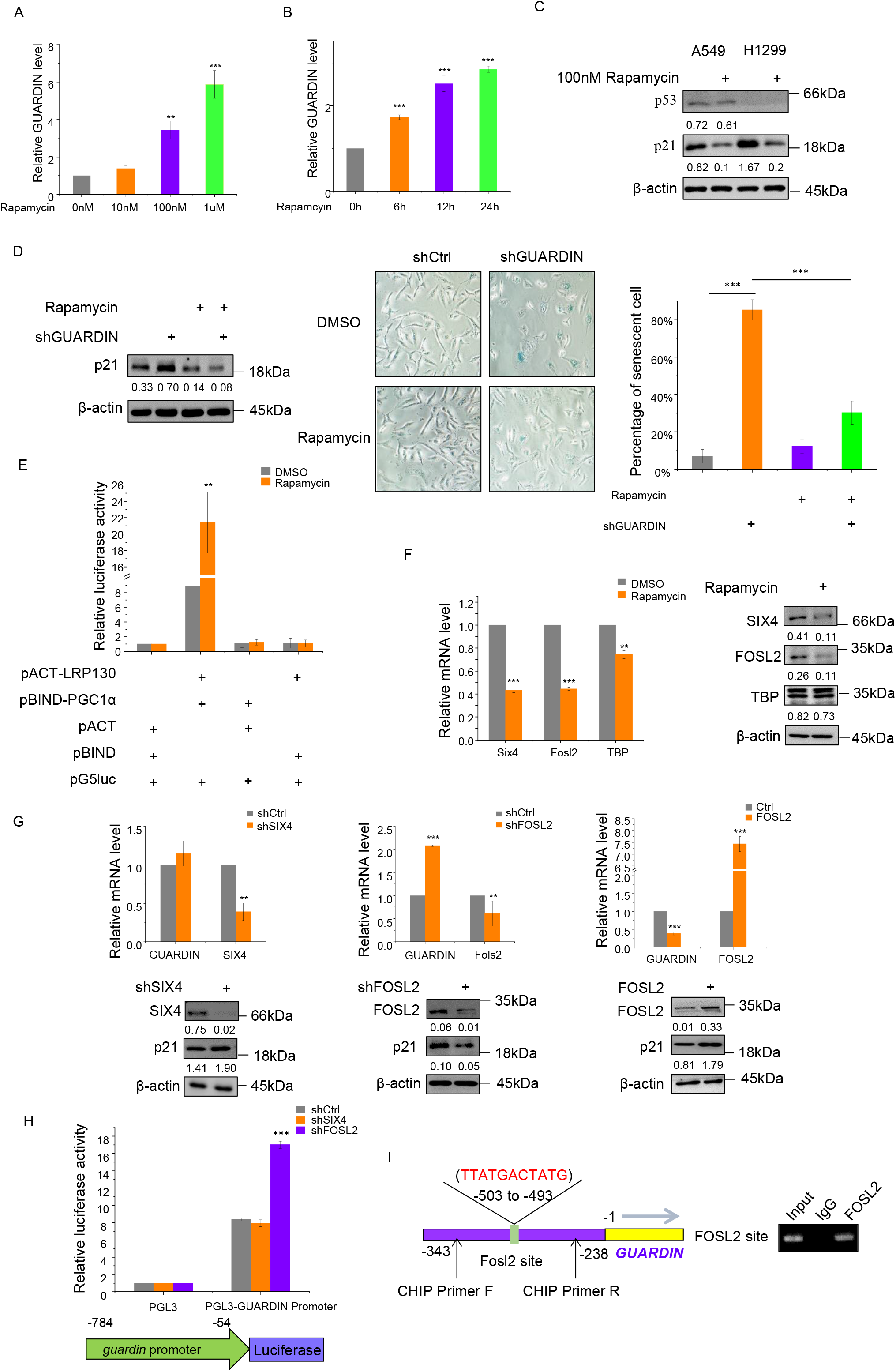
Rapamycin regulates GUARDIN expression via FOSL2. **A, B**. qPCR assays for GUARDIN expression in A549 cells treated with increasing doses rapamycin for 24h (A) or 100nM of rapamycin for the indicated times (B). **C.** Western blotting assays for p53 and p21 expression in A459 and H1299 cells treated with 100nM of rapamycin or vehicle (DMSO) for 48h. **D.** A549 cells with shCtrl or shGUARDIN were treated with 100nM rapamycin or DMSO vehicle in the indicated combinations for 48h. Western blotting was used to measure p21 levels (left) while conducting SA-β-gal staining in parallel (middle) with the percentage of senescent cells calculated from the SA-β-gal staining (right). **E.** Mammalian two-hybrid assays between pACT-LRP130 and pBIND-PGC1α in A549 cells treated with DMSO or 100nM rapamycin. Samples were subjected to the luciferase activity assays. **F.** SIX4 and FOSL2 mRNA (left) and protein levels (right) measured by qPCR and Western blotting, respectively in A549 cells treated with DMSO or 100nM rapamycin. TBP served as a control. **G.** qPCR (upper) and Western blotting (lower) assays for GUARDIN, SIX4 and FOSL2 expression in A549 cells comparing shCtrl with shSIX4 (left), shFOSL2 (middle) or FOSL2 (right) after transduction for 48h. **H.** Luciferase assays in A549 cells transduced with shCtrl, shSIX4 or shFOSL2 using pGL3 (negative control) or pGL3-FOXO4 promoter reporter plasmids. **I.** Schematic illustrating the putative FOSL2 binding site within the *GUARDIN* promoter and the location of primers for ChIP assay (left). RIP assays comparing control IgG versus FOSL2 antibodies demonstrate specific recovery of the GUARDIN promoter by RT-PCR assay (right). (A-G) values are mean± s.e.m (n = 3). (E, F, G) two-tailed paired Student’s t test; (A, B, H) one-way ANOVA with Tukey’s multiple comparison post-test; (D) two-way ANOVA with Bonferroni’s multiple comparison post-test.

To explore the mechanism responsible for the rapamycin-induced upregulation of GUARDIN, we interrogated microarray gene expression data derived from tumor tissues with or without rapamycin treatment (GSE39694; Gene Expression Omnibus (GEO)). Among the 629 genes altered by treatment with rapamycin there were 38 encoding transcription factors (Figure EV 4A, Dataset EV3). Inspecting the genomic sequence in the GUARDIN promoter using the JASPAR database (30) identified putative binding sites for SIX Homeobox 4 (SIX4), FOS-Like Antigen 2 (FOSL2) and ATA-binding protein (TBP) (Figure EV 4B). Further assessment of the levels of these transcription factors showed that SIX4 and FOSL2 were downregulated at mRNA and protein levels, whereas TBP remained unchanged in response to rapamycin treatment (Figure 4F). Moreover, knockdown of FOSL2 but not SIX4 caused upregulation of GUARDIN and downregulation of p21 (Figure 4G), suggesting that FOSL2 expression underpinned the rapamycin-dependent changes in GUARDIN expression. Consistently, overexpression of FOSL2 caused downregulation of GUARDIN and upregulation of p21 (Figure 4G), recapitulating the effects of rapamycin treatment on GUARDIN and p21 expression (Figure 4A, B, C). Additionally, FOSL2 knockdown increased the transcriptional activity of the −784/−54 GUARDIN reporter construct whereas reporter levels after SIX4 shRNA were unchanged from controls (Figure 4H). In addition, ChIP assays demonstrated that FOSL2 bound to its predicted binding sites (−503 to −493) in the GUARDIN promoter (Figure 4I). Together these results indicate that binding of FOSL2 to the GUARDIN promoter transcriptionally represses its expression. Thus, rapamycin suppresses FOSL2 and subsequently upregulates GUARDIN leading to suppression of p21 and inhibition of cellular senescence.

## Discussion

We have previously shown that GUARDIN is a p53-responsive lncRNA and is essential for cell survival and proliferation through maintaining genomic integrity in response to DNA damage (19). In this report, we provide evidence that GUARDIN is also responsive to rapamycin to suppress cellular senescence through facilitating LRP130/PGC1α-mediated repression of FOXO4 leading to transcriptional inactivation of p21. Moreover, we show that repression of FOSL2 is responsible for upregulation of GUARDIN in response to rapamycin. These results not only reveal a p53-independent mechanism that regulates GUARDIN expression, but also uncover a novel signaling pathway of cellular senescence induced by rapamycin.

While replicative exhaustion resulting from telomere shortening or uncapping is a common driver of cellular senescence (21), an important mechanism through which GUARDIN protects genomic integrity is to maintain the expression of telomeric repeat binding factor 2 (TRF2), a component of the shelterin complex critical for capping telomeres (31). Indeed, we found that senescence induced by GUARDIN knockdown was mediated by p21, as knockdown of GUARDIN upregulated p21 and co-knockdown of p21 diminished senescence resulting from GUARDIN knockdown, consistent with the role of GUARDIN in protecting genomic integrity (19). Intriguingly, GUARDIN-mediated regulation of p21 expression appeared to be independent of p53, although p21 is a major effector of p53 signaling and we have previously shown that GUARDIN, similar to p21, may represent an indicator of wild-type p53 activity (19). Nevertheless, p53-independent regulation of p21 has been widely documented and GUARDIN was also detectable in TP53-null cells and tumors with mutations in TP53 (19). Our results further consolidate that notion that both GUARDIN and p21 can function independently of p53 and point to a GUARDIN-mediated mechanism that transcriptionally represses p21 irrespective of p53 activation status.

Previously we have found that GUARDIN acts as an RNA scaffold to facilitate the binding between BRCA1 and BARD in response to DNA damage (19). Intriguingly the primary contribution of GUARDIN in suppressing senescence also involved a role in facilitating protein complexes, in this case the assembly and stability of a repressor complex formed by LRP130 and PGC1α. This was demonstrated by 1), GUARDIN, LRP130 and PGC1α formed a ternary structure; 2), knockdown of GUARDIN reduced the association between LRP130 and PGC1α; 3), binding of LRP130 and PGC1α to the *FOXO4* promoter was associated with transcriptional repression while inhibiting LRP130, PGC1α or GUARDIN increased its activation; and 4), knockdown of LRP130 or PGC1α upregulated p21, recapitulating the effects of knockdown of GUARDIN. Notably the binding interactions between BRCA1/BARD and LRP130/PGC1α were mediated through different regions of GUARDIN (19), indicative of distinct structural specificities embedded in the RNA sequence.

PGC1α is a transcriptional cofactor that interacts with multiple transcription factors (32) while LRP130 is a leucine-rich repeat-containing protein (also known as LRPPRC) functionally associated with mitophagy induction (33) but also with demonstrated transcriptional and/or translational roles in the nucleus and endoplasmic reticulum (34). The LRP130/PGC1α complex plays a critical role in glucose homeostasis and energy balance through regulation of gluconeogenic and mitochondrial genes (25, 35).

Interestingly, previous studies show that PGC1α□is a binding partner and coactivator of FOXO1 which drives the expression of gluconeogenic genes in liver (25, 36) while in contrast, PGC1α acts at the FOXO3 promoter to suppress its expression in skeletal muscle (37). Ours results showing that the GUARDIN/LRP130/PGC1α complex acts to transcriptionally repress FOXO4 are consistent with the latter mode of regulation. Maintaining low FOXO4 levels prevents activation and upregulation of p21 expression and protects cells against senescence induction. Indeed, FOXO4 is known to play an important role in p21 transcriptional regulation (38) and notably, administration of a small interfering FOXO4 peptide reduces senescence and restores overall fitness in naturally aged mice or in fast aging mice models (39). Our results showing suppression of p21 by GUARDIN through LRP130/PGC1α-mediated transcriptional repression of FOXO4 as an important mechanism to counteract cellular senescence implicates that promotion of GUARDIN expression may be a useful approach to prevent aging through sustained repression of FOXO4.

Finally, we establish that GUARDIN expression is induced in cells in response to the mTOR inhibitor, rapamycin. This agent has long been known to inhibit cellular senescence and prolong life span in various model systems (13, 40). However, the mechanisms involved remain less understood and are thought to be multifactorial. We found that GUARDIN contributes to rapamycin-mediated inhibition of cellular senescence, opening a novel scenario that rapamycin protects against cellular aging through a noncoding mechanism. Instructively, treatment with rapamycin did not activate p53, consistent with senescence representing a p53-independent aspect of GUARDIN function. The upregulation of GUARDIN by rapamycin was mediated through downregulation of FOSL2. FOSL2 was shown to exert transcriptional repression of GUARDIN through occupancy of the *GUARDIN* promoter although how rapamycin effects downregulation of FOSL2 is not currently known.

In summary, we have identified a signaling axis encompassing GUARDIN, LRP130/ PGC1α, FOXO4 and p21 that can suppress cellular senescence (Figure 5). Manipulating GUARDIN levels by inhibition or overexpression acted to either trigger or block senescence. An important question involves how the contextual placement of GUARDIN’s functions involving genomic maintenance versus its role in senescence. The latter involves negative control of gene expression that conceivably may act as a failsafe switch under conditions where GUARDIN expression is compromised although this thesis remains to be explored. Nevertheless, as GUARDIN is responsive to treatment with rapamycin, our findings also highlight a potential role for GUARDIN in aging-associated diseases with perhaps practical implications in aging-prevention.

**Figure 5.**
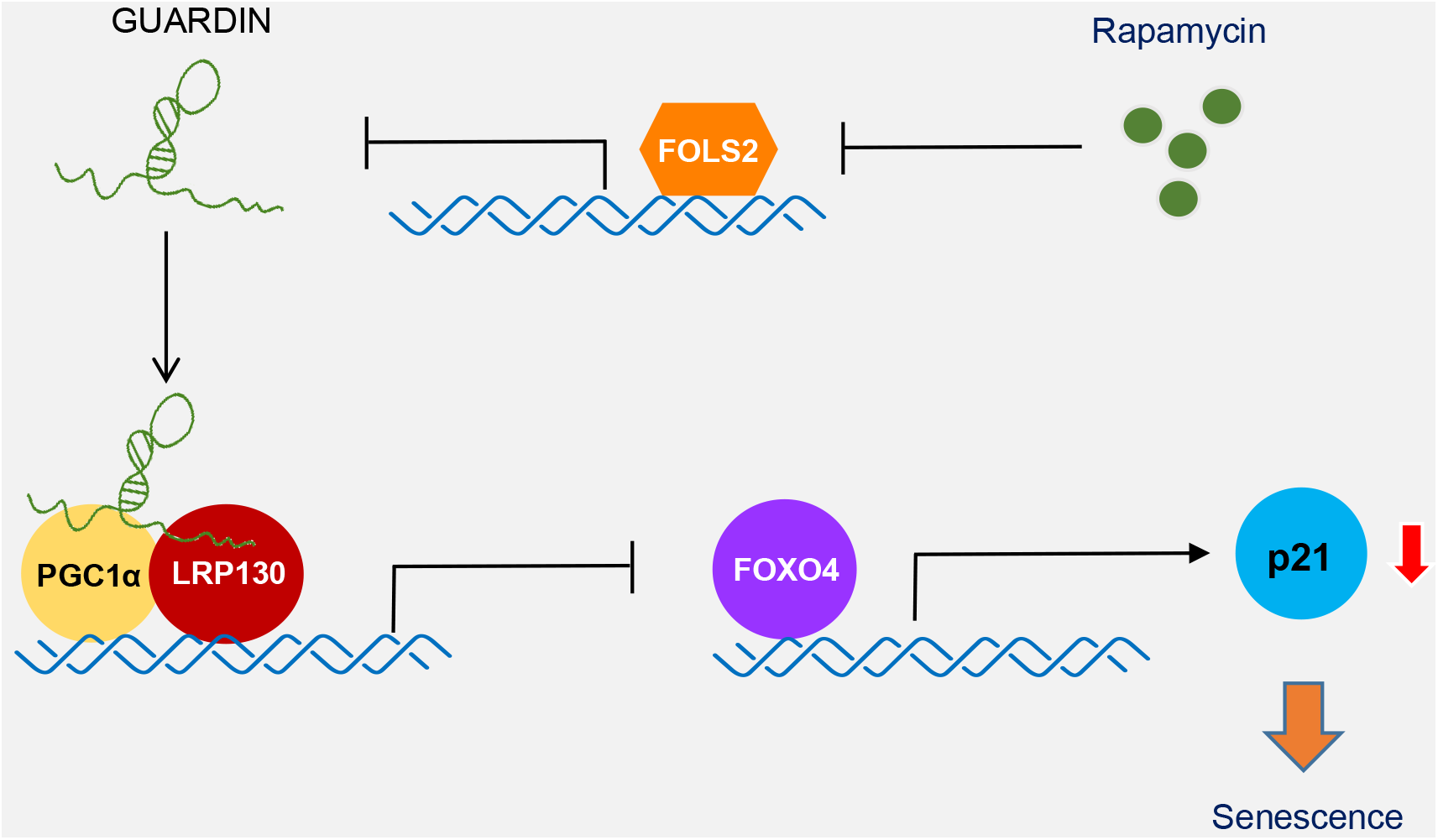
Model for GUARDIN-mediated regulation of cellular senescence. GUARDIN acts as an RNA scaffold to facilitate interaction between LRPl30/PGC1α that acts to transcriptionally repress FOXO4 expression. This repression prevents FOXO4-mediated upregulation of p21 and cell entry into senescence. GUARDIN participates in rapamycin-induced cellular senescence, likely through downregulation by FOLS2, which is also downregulated by rapamycin.

## Materials and Methods

### Reagents and antibodies

Reagents and antibodies were sourced as indicated: cycloheximide (Sigma); rapamycin (BBI); actinomycin D (Solarbio); streptavidin beads (Invitrogen); A/G beads (Pierce); anti-H3K9me3, p15, p16 and Alexa Fluor^®^ 488-conjugated secondary antibodies (Abcam); anti-p21, flag (Sigma); anti-IL6, IL-8, LRP130, HSPA5, MPG, SIX4, FOLS2, TBP and FOXO4 (Proteintech); anti-GAPDH and β-actin (CMC-TAG); anti-HUR, H2A, p53, ROD1 and PGC1α (Santa Cruz); HRP-conjugated secondary antibodies against mouse and rabbit IgG (Promega).

### Cell culture

HepG2, A549, H1299, 293T and HAFF cells were cultured in DMEM (Invitrogen) supplemented with 10% FBS / 1% penicillin/streptomycin and maintained in a humidified incubator at 37 °C/ 5% CO2. All cells were authenticated by STR profiling (GenePrint 10 System kit from Promega and AuthentiFiler PCR Amplification Kit from ThermoFisher) and tested negative for mycoplasma contamination using the Cell Culture Contamination Detection Kit (ThermoFisher).

### RNA interference and gene overexpression

Lentiviruses for gene knockdown and overexpression experiments, respectively, were generated by transfecting HEK293T cells for 48 h with PLKO.1 based shRNAs, pREV, pGag and pVSVG (2:2:2:1 ratio) or psin/pCDH, pspax2, pmd2.g (2:2:1 ratio). Supernatants were 0.45 um filtered, supplemented with 8 μg/ml polybrene (Sigma) before incubating with target cells and selecting with 1 mg/ml puromycin. Targeting sequences are shown in Appendix Table S1.

### Quantitative and semi-quantitative RT-PCR

Total RNA was isolated by TRIzol reagent (Invitrogen) and 500 ng RNA used to synthesize cDNA using PrimeScriptTM RT reagent kit (TaKaRa) according to the manufacturer’s instructions. Semi-quantitative RT-PCR was performed using 2x Taq PCR mix with 25 cycles for internal controls and 30-40 cycles for RNAs. Primer sequences are shown in Appendix Table S1. QPCR was performed as described previously (41).

### β-Galactosidase staining and quantitation

β-galactosidase staining was performed according to the manufacturers protocol (C0602, Beyotime). β-galactosidase-positive cells were scored using light microscopy from three random fields.

### Immunofluorescence and RNA FISH

Cells grown on coverslips were fixed in 4% paraformaldehyde and permeabilized using 0.2% Triton X-100. After blocking with 2% BSA, coverslips were incubated overnight at 4 °C with primary antibodies before addition of Alexa-488 secondary antibodies. RNA probes synthesized by T7-mediated transcription were purified by phenol-chloroform extraction and 1 mg RNA labelled with Alexa-546 using the ULYSIS Nucleic Acid Labeling Kit (Thermo Fisher (42). Nuclei were counterstained with Hoechst.

### ELISA

Cells were incubated in serum-free DMEM for 24 h and IL-6/8 levels determined in supernatants using the femtoELISA™ HRP Kit (G-Biosciences).

### Western Blotting

Whole cell lysates were prepared using RIPA buffer containing protease inhibitors (Beyotime) before conducting SDS-PAGE and Western blotting using enhanced chemiluminescence.

### Northern Blotting

Northern blots were performed as described previously (41) against total RNA resolved on 1.5% agarose gels. Membranes were hybridized with digoxigenin-labeled antisense GUARDIN probes synthesized using T7 RNA polymerase using the DIG Northern Starter Kit (Roche).

### Biotin pull-down assays and mass spectrometry

Biotinylated sense (negative control) and antisense biotin-labeled DNA oligomers corresponding to GUARDIN (1μg) were coupled to streptavidin-coupled Dynabeads (Invitrogen). The Dynabeads were subsequently incubated with cell lysates for 4 h and eluted proteins subjected to SDS–PAGE and staining with Coomassie Brilliant Blue G-250. Protein bands were excised and sent to Core Facility of Center for Life Sciences, USTC for mass spectrometry (MS) analysis using a Thermo-Finnigan LTQ LC/MS-MS. Proteins IDs determined by MS are listed in Dataset EV2. All processes were performed under RNase-free conditions.

### Immunoprecipitation

Cells were lysed in IP lysis buffer (150 mM NaCl, 50 mM Tris, pH 7.4, 10% glycerol and 1.5 mM MgCl_2_) supplemented with protease inhibitor cocktail before incubation for 4 h at 4°C with protein A/G beads precoated with indicated antibodies. Beads were washed three times using IP lysis buffer, eluted with heat (95°C, 10 min) and the samples analyzed by Western blotting. Where indicated, two step IPs were performed against cell lysates prepared using lysis buffer containing 20 mM HEPES, pH 7.8, 400 mM KCl, 5% glycerol, 5 mM EDTA, 1% NP40, protease inhibitors cocktail and RNase inhibitor. The first IP used anti-Flag antibodies before elution with Flag peptides. Ten percent of the sample was reserved for Western blotting and semi-quantitative RT–PCR analysis, respectively, while the remaining eluate was subjected to the secondary IP. Steps were performed under RNase free conditions as required.

### RNA immunoprecipitation

RNA immunoprecipitation (RIP) was performed as described (43). Briefly, cells were lysed in RIP buffer supplemented with RNase A inhibitor and DNase I. Soluble cell lysates were precleared with protein A/G beads alone before IP with the indicated antibodies at 4°C for 3 hr. After washing, the bead-bound immunocomplexes were eluted using elution buffer (50 mM Tris, pH 8.0, 1% SDS, and 10 mM EDTA) at 65°C for 10 min. To isolate protein-associated RNAs from the eluted immunocomplexes, samples were treated with proteinase K, and RNAs were extracted by phenol/chloroform before RT-PCR analysis.

### In vitro transcription

The T7 RNA polymerase promoter sequence was introduced into DNA templates generated by PCR to allow for in vitro transcription. PCR products were purified using the DNA Gel Extraction Kit (Axygen), and in vitro transcription performed using the T7-Flash BiotinRNA Transcription Kit (Epicentre, biotin labelling) or TranscriptAid T7 High Yield Transcription Kit (ThermoScientific, unlabelled) according to the manufacturer’s instructions. RNA was subsequently extracted using phenol-chloroform. Primer sequences are shown in Appendix Table S1.

### ChIP assay

Cells were crosslinked with 1% formaldehyde for 10 min and ChIP assays performed using the Pierce Agarose ChIP kit (ThermoScientific, USA) according to the manufacturer’s instructions. Anti-rabbit immunoglobulin G was used as a negative control. Bound DNA fragments were subjected to RT-PCR using the specific primers (Appendix Table S1).

### Luciferase reporter assay

Luciferase reporter assays were performed as described previously (43) using the Dual-Luciferase Reporter Assay System (Promega, Madison, WI, USA).

### Subcellular fractionation

Cells were incubated with hypotonic buffer (25 mM Tris-HCl, pH 7.4, 1 mM MgCl_2_, 2.5 mM KCl) on ice for 5 min. An equal volume of hypotonic buffer containing 1% NP-40 was then added, and each sample was left on ice for another 5 min. After centrifugation at 5000 g for 5 min, the supernatant was collected as the cytosolic fraction. The pellets were re-suspended in nuclear resuspension buffer (20 mM HEPES, pH 7.9, 400 mM NaCl, 1 mM EDTA, 1 mM EGTA, 1 mM DTT, 1 mM PMSF), and incubated at 4°C for 30 min. Nuclear fractions were collected after removing insoluble membrane debris by centrifugation at 12,000 g for 10 min.

### RNA-seq

Total RNA was extracted by phenol/chloroform and RNA-seq analysis performed by Shanghai Biotechnology.

### Mammalian Two-Hybrid assays

Complementary DNAs for PGC1α and LRP130 cloned into the pBIND and pACT vectors were transfected along with pG5luc vector (Promega) and reporter activity measured 48 h later using the Dual-Luciferase Reporter Assay Kit (Promega). Renila measurements were used to normalize changes in firefly luciferase activity.

### Quantification and Statistical analysis

Statistical analysis was carried out using Microsoft Excel 2016 and GraphPad Prism 7 to assess differences between experimental groups. Densitometry was performed using Image-Pro plus 6.0 software. Statistical significance was analyzed by two-tailed Student’s t-test for comparisons of two samples, one-way ANOVA with Tukey’s post-test for univariate comparisons, two-way ANOVA with Bonferroni’s post-test for bivariate comparisons. P values lower than 0.05 were considered to be statistically significant. (ns, not significant, * P < 0.05, ** P< 0.01, *** P < 0.001).

### Data Availability

Raw and normalized data files for the microarray analysis have been deposited in the NCBI Gene Expression Omnibus under accession number GSE39694.

## Acknowledgements

Authors thank Dr. Qidong Li for valuable discussions and technical support. We also thank Drs. An Xu and Wanglai Hu for providing full-length GUARDIN constructs. This work was supported by grants from National Key R&D Program of China (2018YFA0107103 and 2016YFC1302302), the National Natural Science Foundation of China (81820108021, 81430065, 31871437, 31601146 and 81772908) and the National Health and Medical Research Council of Australia (1147271).

## Author contributions

XDS, SSF, XYL and MW designed the research. XDS and MH performed the experiments and data analysis. XDZ and RFT participated in the data analysis. XDZ, RFT, XYL and MW wrote the manuscript.

---

**Figure EV 1.**
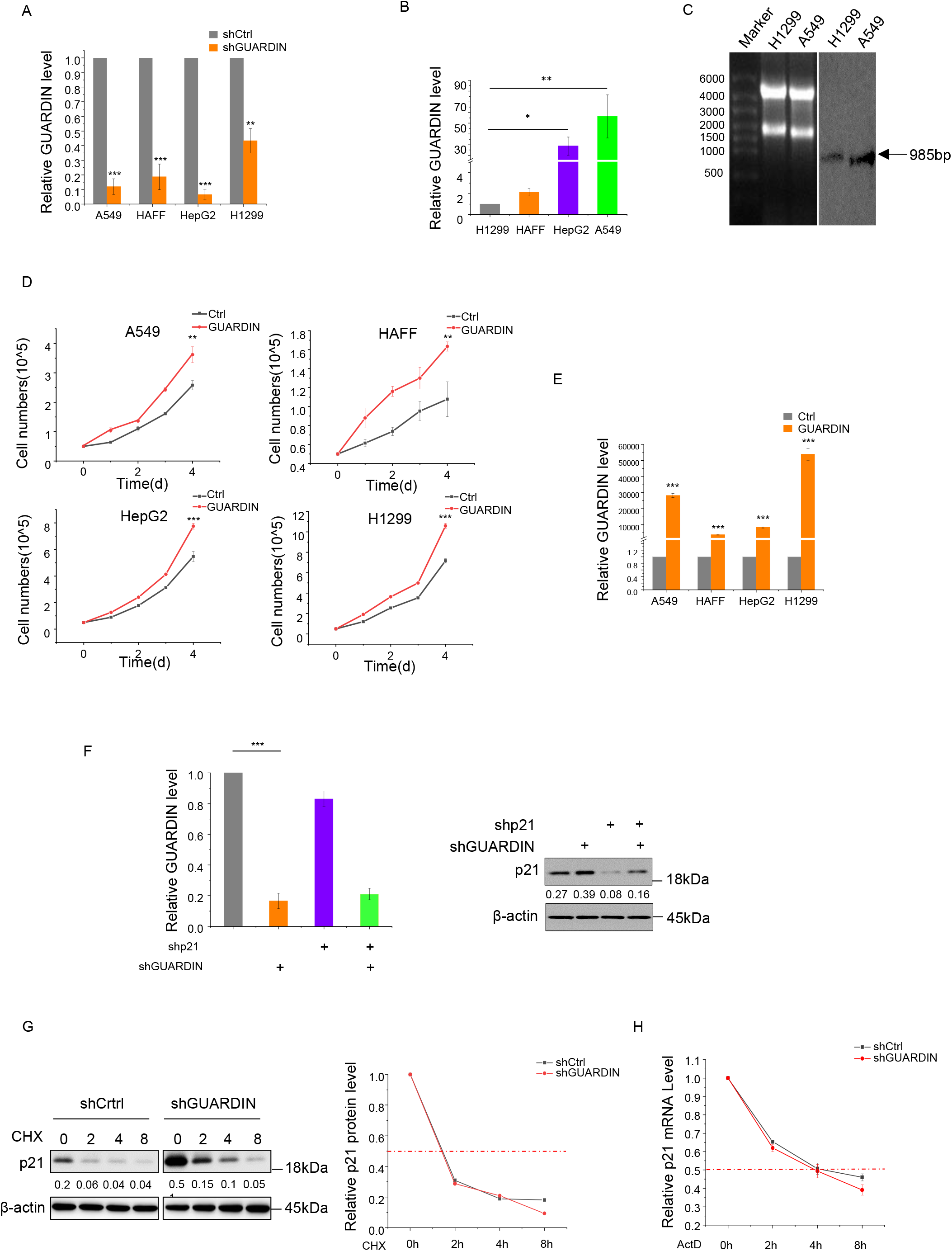
**A.** shRNA knockdown efficiency. GUARDIN levels in A549, HAFF, HepG2 and H1299 cells were measured using qPCR. **B. C:** lncRNA validation. B. Comparative levels of GUARDIN in A549, HAFF, HepG2 and H1299 cells measured using qPCR. All data normalized to H1299 levels. **C.** Northern blot against GUARDIN using total RNA from H1299 and A549 cells. 28S and 18S RNA loading controls (left) and GUARDIN (right) resolving as a single band. **D.** Overexpression of GUARDIN promotes cell proliferation *in vitro.* Cell numbers were counted in A549, HAFF, HepG2 and H1299 cells with and without GUARDIN overexpression for 4 days (d). **E.** Overexpression efficiencies. qPCR analysis of GUARDIN level in A549, HAFF, HepG2 and H1299 cells after 24h transduction with GUARDIN. **F.** Co-silencing of GUARDIN and/or p21. qPCR assays for GUARDIN (left) and Western blotting for p21/actin (right) in A549 cells after 24h transduction with shCtrl or shGUARDIN. **G. H.** Silencing of GUARDIN does not affect p21 protein or mRNA stability. G. A549 cells were treated with 50 ng/ml of cycloheximide (CHX) for the indicated times before measuring p21/□□ Lactin by Western blotting (left). Densitometric analysis (right) of relative p21 levels. H. p21 mRNA levels were measured by qPCR in A549 cells treated with 1 mg/ml of actinomycin D (actD) for the indicated time points. (A, B, D, E, F, H) values are mean± s.e.m (n= 3). (A, D, E, H) two-tailed paired Student’s t test; (B) one-way ANOVA with Tukey’s multiple comparison post-test; (F) two-way ANOVA with Bonferroni’s multiple comparison post-test.

**Figure EV 2.**
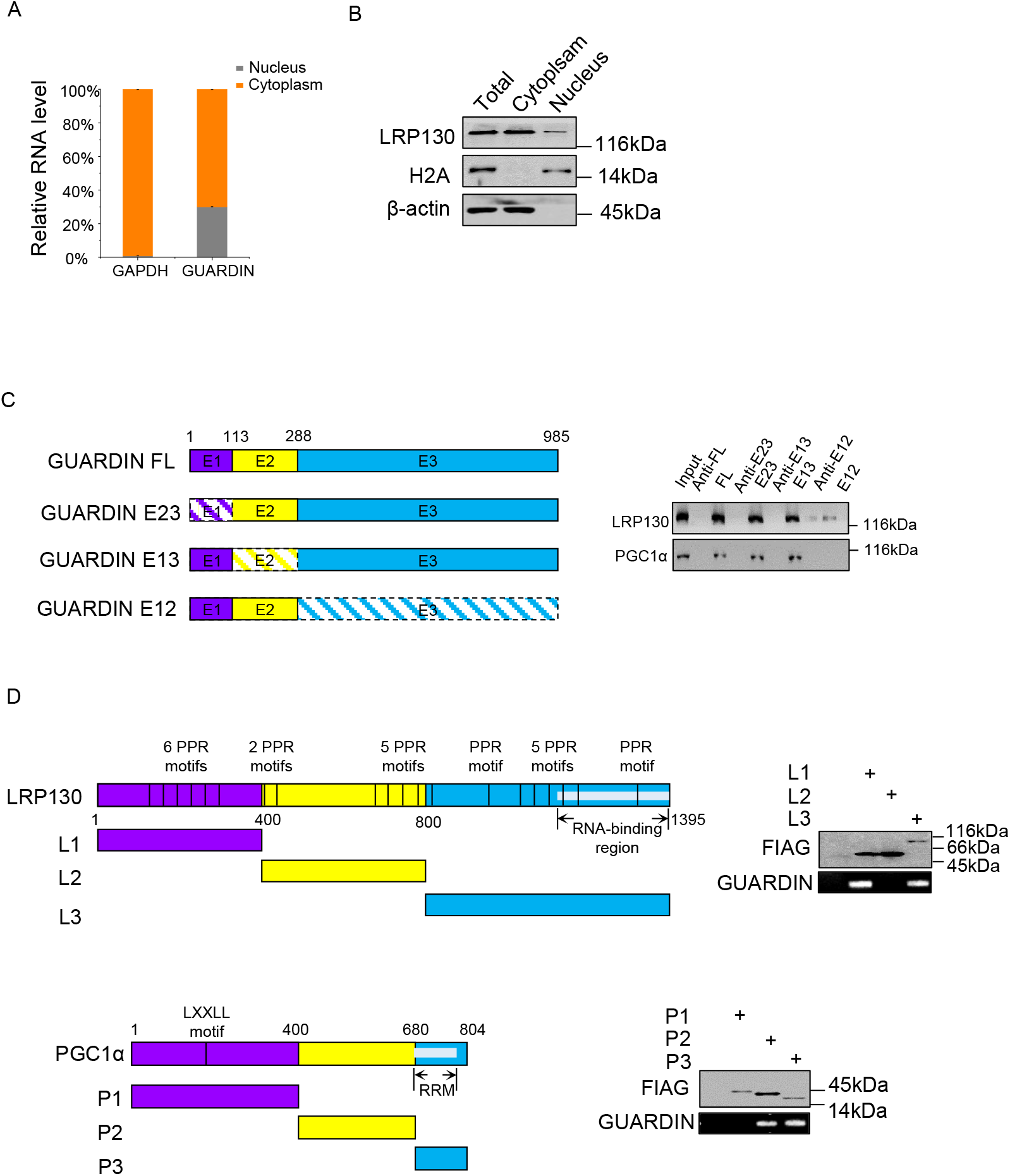
**A, B.** Subcellular distribution of GUARDIN and LRP130. A. RT-PCR assays against GUARDIN in cytoplasmic and nuclear compartments isolated from A549 cells. GAPDH served as cytoplasmic control. B. Western blotting against LRP130 in distribution in subcellular fractions from A. H2A and β-actin served as nuclear and cytoplasmic markers, respectively. **C.** GUARDIN interacts with LRP130/PGC1α through exon3 (E3). Schematic illustration of GUARDIN and the design of exon-deletion constructs (left). RNA pulldown assays against whole-cell lysates of A549 cells using in vitro-transcribed GUARDIN or mutant RNAs (right). Western blotting assay was performed for LRP130 and PGC1α. **D.** LRP130 interacts with GUARDIN by its N-terminal and C-terminal domains (upper), while PGC1α interacts with GUARDIN by its C-terminal and central region domains (lower). Schematic illustration of LRP130 and PGC1α and the design of domain truncation constructs (left). RIP assays performed against whole-cell lysates of A549 cells transfected with indicated FLAG-tagged LRP130/PGC1α mutant constructs (left). Western blotting assays for LRP130/PGC1α mutants using FLAG antibodies and RT-PCR assays for GUARDIN (right). (A) values are mean± s.e.m (n= 3).

**Figure EV 3.**
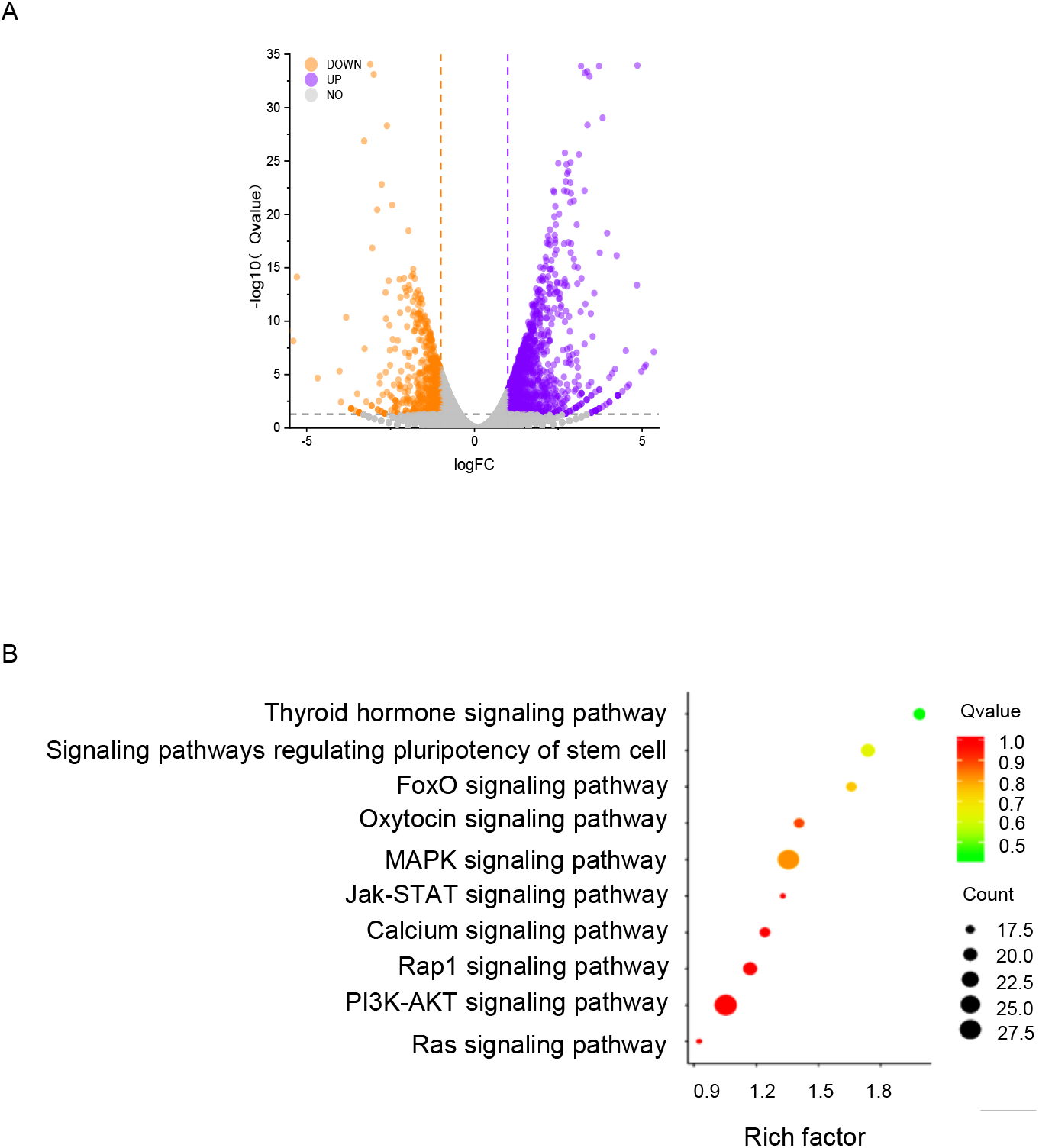
**A.** Differentially expressed mRNAs in A549 cells following shGUARDIN knockdown. Volcano plot comparing control versus GUARDIN knockdown in A549 cells with differentially downregulated and upregulated mRNAs shown in orange and violet, respectively. **B.** KEGG pathway analysis. Selected high scoring KEGG pathway enrichments using data from (A) are illustrated in the y-axis. Q values (downregulated or upregulated) pathways are represented by color with the size of the dot indicating the number of DEGs mapped to each pathway.

**Figure EV 4.**
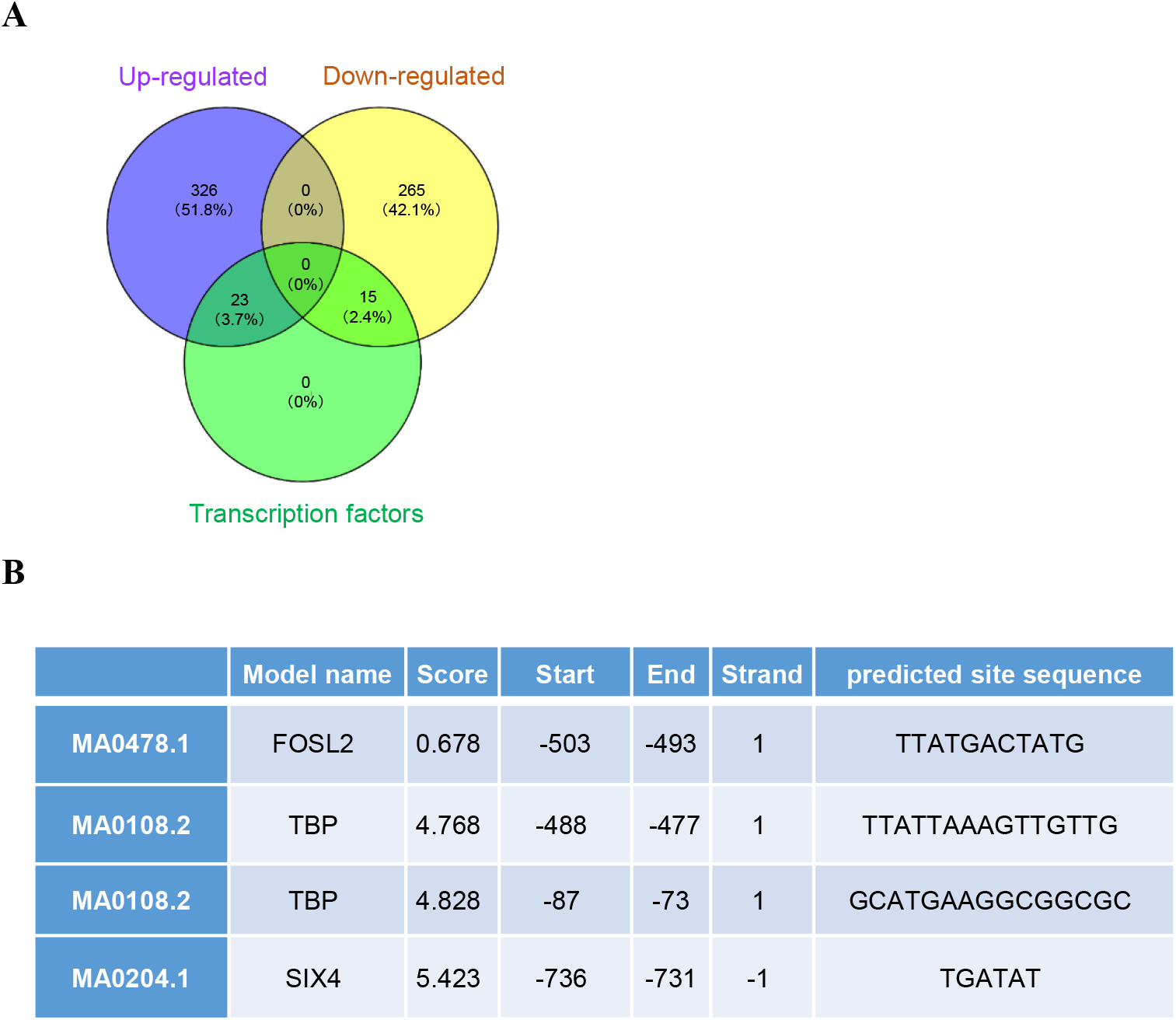
**A.** DEGS in tumor tissues treated with rapamycin versus control groups determined by RNA-seq. Venn diagram shows upregulated transcripts (violet), downregulated transcripts (yellow) along with those identified as transcription factors (green). **B.** Transcription factors consensus binding sites present within the *GUARDIN* promoter.

